# Self-rectifying magnetoelectric metamaterials enable precisely timed remote neural stimulation and restoration of sensory motor functions

**DOI:** 10.1101/2022.01.24.477527

**Authors:** Joshua C. Chen, Gauri Bhave, Fatima Alrashdan, Abdeali Dhuliyawalla, Jacob T. Robinson

**Author notes:** These authors contributed equally to this work (Joshua C. Chen, Gauri Bhave).

## Abstract

Magnetoelectric materials convert magnetic fields to electric fields and have applications in wireless data and power transmission, electronics, sensing, data storage, and biomedical technology. For example, magnetoelectrics could enable precisely timed remote stimulation of neural tissue, but the resonance frequencies where magnetoelectric effects are maximized are typically too high to stimulate neural activity. To overcome this challenge, we created the first self-rectifying magnetoelectric “metamaterial.” This metamaterial relies on nonlinear charge transport across semiconductor layers that allow the material to generate a steady bias voltage in the presence of an alternating magnetic field. This “self-rectification” allows us to generate arbitrary electrical pulse sequences that have a time-averaged voltage in excess of 1 V. As a result, we can use magnetoelectric nonlinear metamaterials (MNMs) to remotely stimulate peripheral nerves with repeatable latencies of less than 5 ms, which is more than 120 times faster than previous neural stimulation approaches based on magnetic materials. These short latencies enable this metamaterial to be used in applications where fast neural signal transduction is necessary such as in sensory or motor neuroprosthetics. As a proof of principle, we show wireless stimulation to restore a sensory reflex in an anesthetized rat model as well as using the MNM to restore signal propagation in a severed nerve. The rational design of nonlinearities in the magnetic-to-electric transduction pathway as described here opens the door to many potential designs of MNMs tailored to applications spanning electronics, biotechnology, and sensing.

---

Remote stimulation of neural activity would enable less invasive medical therapies and improve our ability to study neuroscience in freely behaving animals. Magnetic fields are a promising candidate for remote control of neural activity because these fields can pass losslessly through air and penetrate deep into the body. However, the cells and proteins in our body have very weak magnetic properties; thus, for magnetic fields to stimulate a biological response, researchers typically use high-field strengths in excess of 1 T that are generated by pulsed magnetics (*1*) or magnetic materials that convert a weak magnetic stimulus (typically less than 100 mT) into a form of energy that can stimulate nearby cells. Indeed, this materials-based approach offers superior selectivity because one can target a specific area for stimulation by limiting the location of the magnetic materials, or one can combine the magnetic materials with genetically modified cells to achieve cell-type specific neural stimulation (*2*–*7*). A materials-based magnetic stimulation technique that could achieve millisecond timing would enable numerous neurotherapeutic applications and research applications that require neuromodulation to be precisely timed with sensory stimuli and behavior (*8*); however, existing magnetic materials have fallen well short of millisecond temporal precision. For example, magnetic heating of superparamagnetic nanoparticles can stimulate neurons that express thermoreceptors (*2*–*5*) with behavioral response times approaching 500 ms (*6*) or trigger the release of drugs that cause a behavioral response with a latency of several seconds (*7*). Mechanical displacement of magnetic nanoparticles can also stimulate neural activity via mechanoreceptors (*9*,*10*), but the response times are several seconds in vivo (*11*).

Magnetoelectric (ME) materials are a promising candidate for temporally precise remote neural stimulation but have yet to demonstrate millisecond timing. Because ME materials convert magnetic fields to electric fields they can, in principle, activate voltage-gated channels to drive precisely timed neural activity just like an electrode. However, large ME coupling coefficients are only found at frequencies too fast to efficiently stimulate neurons. These large ME coupling coefficients are found in strain-mediated ME materials that consist of piezoelectric and magnetostrictive materials. The conversion between magnetic and electric field is maximized when the applied magnetic field matches a mechanical resonant mode of the material (*12*). For materials less than 5 mm in length the lowest resonance frequency is typically larger than 100 kHz which is too fast to directly stimulate neural activity (*13*). As a result, ME materials have only enabled remote neural stimulation when they are driven off resonance where the weak ME coupling leads to a seconds-long latency in the neural response or requires extremely high applied magnetic field strengths to reduce the latency to a few hundred milliseconds (*14*,*15*). Precisely timed neural stimulation with millisecond latencies has only been achieved using ME materials when they are combined with electronic circuits that convert the high-frequency ME response to a low-frequency electrical stimulus (*16*–*19*).

Here we demonstrate targeted remote stimulation of neural activity with millisecond latency by creating the first self-rectifying magnetoelectric *metamaterial* (Fig. 1A). A self-rectifying material converts an alternating current (AC) driving signal to a direct current (DC), or steady bias voltage. This process allows us to generate a low frequency voltage pulse by simply pulsing an AC carrier wave (Fig. 1A). While self-rectification has been observed in many systems with a large second-order nonlinearity, it has yet to be reported in ME materials even under conditions with a large second-order nonlinearity. The lack of self-rectification in ME materials has been explained by a finite conductance across ME laminates that quickly screens any bias that develops across the material (*20*,*21*)

**Fig 1.**
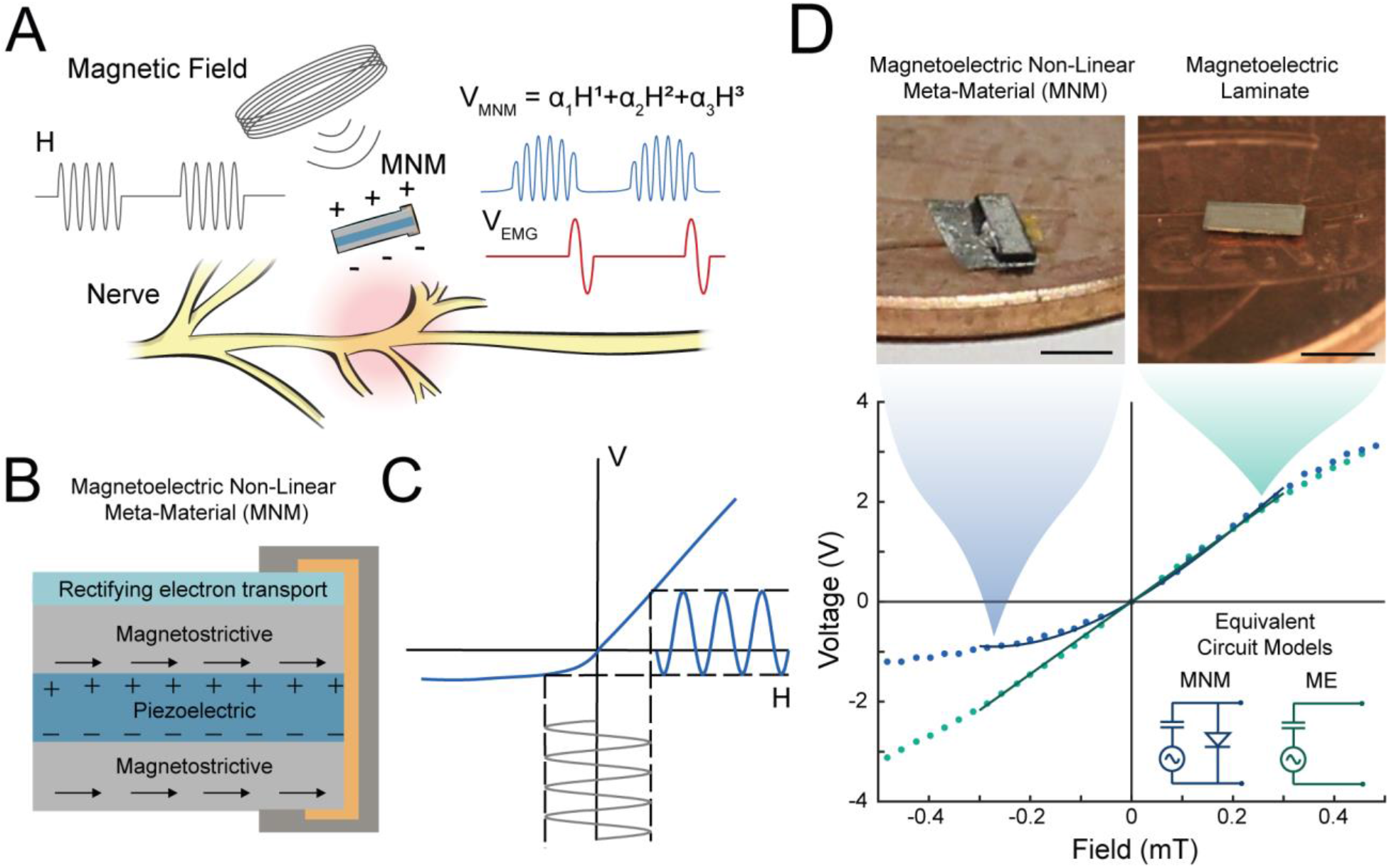
(A) Schematic of the remote neural stimulation using a magnetoelectric metamaterial (MNM). Upon applying a pulsed magnetic field, the MNM generates a biased non-linear electric field that is used to stimulate neural tissue. (B) Schematic of the laminate structure of the MNM. The metamaterial consists of a piezoelectric layer between two magnetostrictive layers. A rectifying electron transport (RET) layer sits on top of the stack and includes an electrical connection between the top and bottom surfaces. This additional layer introduces a non-linear Magnetoelectric (ME) coupling coefficient as seen when plotting the relationship between H and V. (C) Conceptual plot of the voltage obtained across the MNM upon application of an AC magnetic field at the mechanical resonant frequency of the material. Note that the nonlinear H-V relationship causes self-rectification of the voltage. (D) Measured magnetic field (H) vs Voltage (V) curves for both the MNM (blue) and ME laminate (green) along with a polynomial and a linear fit, respectively. Inset shows the two equivalent circuit models for both the MNM and ME and magnified images of each material is displayed on top (Scale bars = 2 mm).

The concept of a self-rectifying ME material is inspired by the field of optical metamaterials, which showed that incorporating sub-wavelength-sized electronic devices into a material allows one to engineer effective dielectric permeabilities with large nonlinear properties (*22*,*23*). We hypothesized that we could use a similar approach to create a self-rectifying magnetoelectric nonlinear metamaterial (MNM). Specifically, we speculated that by adding a nonlinear charge transport layer to the ME laminate (Fig 1B) we could engineer a nonlinear magnetoelectric coupling coefficient α that describes the relationship between the applied magnetic field (H) and the induced electric potential (V) (V = α H) (Fig 1C). For strain-mediated ME materials α is the result of 4 separate processes: 1) Magnetic field (H) applied to a magnetostrictive phase of the material creates stress 2) Stress is transferred into a piezoelectric layer of the material to induce strain 3) Strain in the piezoelectric layer generates an electrical displacement (D), and 4) the Electrical displacement generates an electrical potential (V) across the film based on the effective dielectric properties of the material. In most strain-mediated ME materials operating at the point of maximum ME coupling, these four energy transfer processes are linear (*24*) (Fig. S1).

To create a self-rectifying MNM we added a nanoscale rectifying electron transport (RET) layer that creates a nonlinear relationship between the electric displacement (D) and the electrical potential (V) (Fig 1D). We started with an ME material composed of a tri-layer laminate with a piezoelectric layer made of PZT at the center of two magnetostrictive layers made of Metglas (*25*) (Fig. S1). These layers are mechanically coupled using a two-part epoxy (see methods). We then added the non-linear RET layer to the ME laminate by depositing 130 nm of ZnO and 40 nm of HfO_2_ to create a thin film stack that acts as a diode for charge carriers moving through the material (*26*,*27*) (Fig. 2D). Our rationale for this design was to break the symmetry in the voltage response to positive and negative magnetic fields. In this way we could create a strongly non-linear ME coupling coefficient that would result in self-rectification (Fig. 1C). We chose ZnO for the RET layer because it has been used previously to fabricate diodes (*28*,*29*) and we can deposit it at low temperatures (100 °C) without the need to perform chemical etching, which allows monolithic integration with ME laminates. To improve the rectifying characteristics of the structure, we controlled the electron concentration in the ZnO layer by using Atomic Layer Deposition (ALD) at a low temperature with regulated precursor flow rates (*30*,*31*). We also deposited a passivating HfO_2_ layer to further improve rectification characteristics by preventing reverse current (Fig 2E) (see methods). This thin-film stack works exclusively in Space Charge Limited Current (SCLC) mode thus overcoming the drawbacks of diodes that rely on tunneling currents (Fig. S2B). Characterization of the ZnO/HfO_2_ stack showed typical diode rectification properties with an ideality factor of 6.8 and a rectification ratio of 95.38 (see Methods, Fig. S2A). With the addition of the RET, we can model the MNM material as a clamper circuit (Fig 1D bottom inset), which shows a nonlinear ME response.

**Fig. 2.**
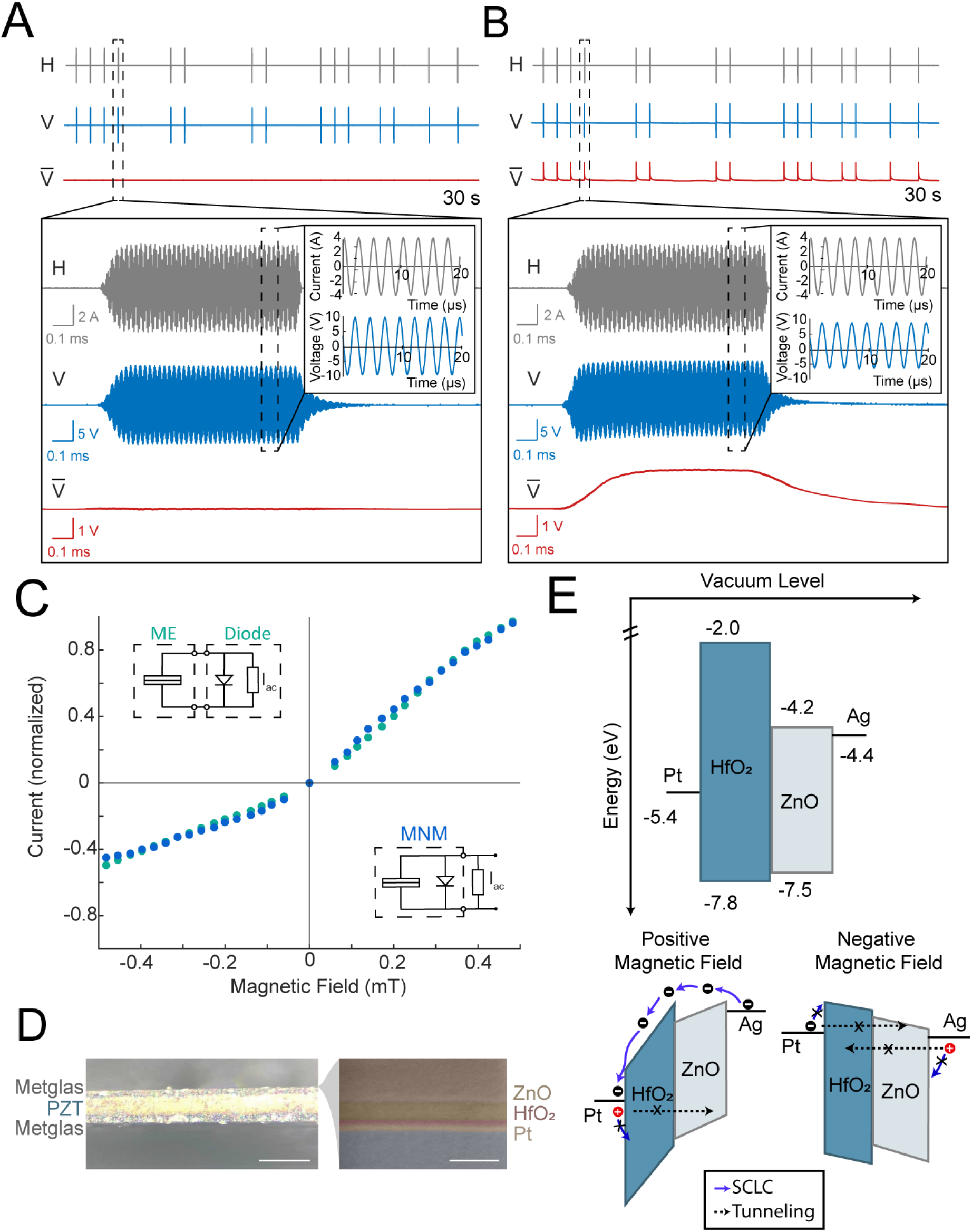
(A & B) Top: Gray lines show arbitrary pulse train of an applied AC magnetic field (H) at the mechanical resonant frequencies of ME and MNM materials, respectively. Blue lines show the measured voltages from ME and MNM materials. The red line shows the voltage resulting from averaging across a 200 μs sliding window of the pulsed voltage waveform. For the MNM we observe a self-rectification of 1.7 V in the average voltage, but for the ME the average voltage remains near zero. Bottom: Magnified time scales of the pulse trains to show a single pulse. Inset shows further magnification revealing the self-rectifying response in the MNM. All data taken at the fundamental longitudinal resonance modes. (C) Normalized magnetic field (H) vs voltage (V) plotted for a ME laminate connected to a RET (green) and for an integrated MNM (blue). The two plots closely follow each other confirming that the nonlinear ME coefficient can be well explained by the altered electronic properties produced by the RET. (D) Optical image of cross section of MNM material. (Scale bar =150 μm) Expanded false-colored SEM image of the RET layers. (Scale bar = 300 nm) (E) Band diagram of the RET layers showing both the forward and reverse biased modes when a positive magnetic field is applied vs a negative field.

When we characterized the ME coupling coefficient of the MNM we indeed found a strong 2nd and 3rd order nonlinearity (Fig. 1D). Plotting the induced voltage across the MNM as a function of the amplitude of the applied magnetic field between −0.5 mT and 0.5 mT (Fig 1D, blue dots) we see a nonlinear relationship that can be well-fit as a third order polynomial, where V =α_1_H +α_2_H_2_ + α_3_H_3_ (α_1_= −7.75, α_2_= 7.75 and α_3_= 5.99) (Fig. 1D, blue line). This is in stark contrast to unmodified ME laminates (Fig 1D, green dots) that are well fit by a linear relationship V=α_1_H (α_1_= 7.2405) (Fig. 1D, green line) (see methods). In all cases these experiments were performed with the alternating magnetic field frequency matching the fundamental resonant mode and under a DC magnetic field bias where the ME coefficient is known to achieve the maximum value (see Methods).

We confirmed that the nonlinear ME coupling coefficient is the result of the nonlinear RET by comparing the MNM response to the nonlinear transport across the RET layer (Fig. 2C). Figure S3A shows the I-V relationship measured from our RET, at the operational frequency of 335 kHz, which is the mechanical resonant frequency of the MNM. We then used the linear H-V relationship measured in the ME laminate (Fig 1D) to convert the I-V curve in Fig S3A, into an H-I curve in Fig S3B (see Methods). We separately measured the H-I relationship of the MNM by varying the amplitude of the alternating magnetic field (H) while measuring the current (I) across a load resistor (R_L_) (Fig. 3C). When we normalized the H-I curve measured in the MNM and computed H-I curve for the ME+RET we find that both H-I relationships show similar nonlinear relationships (Fig. 2C). These results confirm that the nonlinearity observed in the MNM can be explained by the non-linear charge transport across the RET layer (Fig. 2C).

**Fig. 3.**
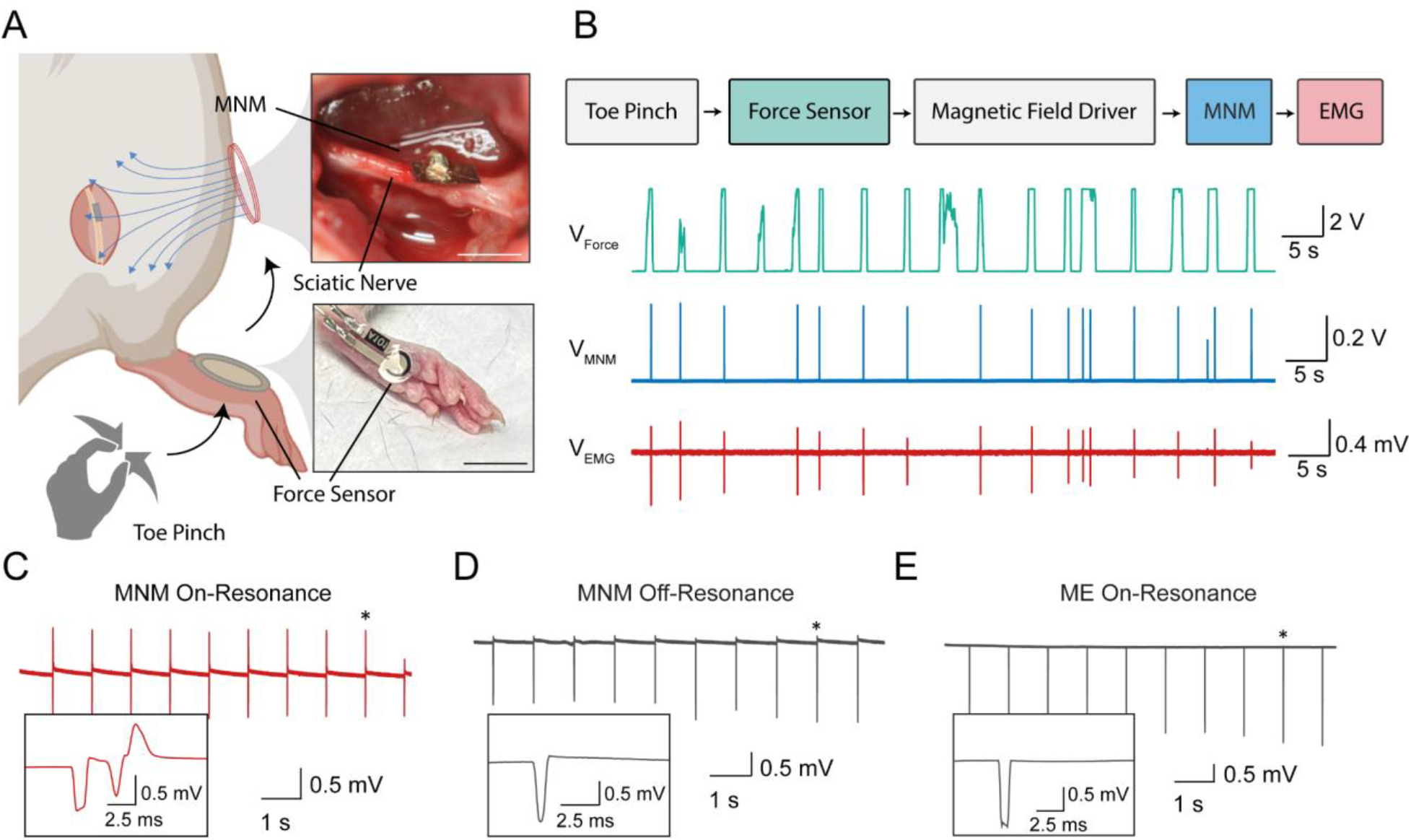
(A) Schematic of sensory and reflex restoration based on a toe pinch in an anesthetized rat. Inset images show a representative MNM in contact with the rat sciatic nerve (scale bar = 3 mm) and the force sensor used on the rat foot (scale bar = .1 cm). (B) Flow chart depicting sensory pathway from toe pinch to resulting leg contraction measured by the EMG. Voltage recordings of the force sensor, MNM, and EMG show that arbitrary patterns of toe pinches produce a restored motor reflex. (C) EMG response measured when MNM is driven with a magnetic field at the fundamental longitudinal resonance frequency and pulsing at a periodic 1 Hz. (D) Control experiment shows no measurable EMG response obtained when MNM is driven off-resonance. (E) Second control experiment shows no measurable EMG response when MNM was replaced with similarly sized ME laminate and driven on resonance.

When we apply an AC magnetic field at the mechanical resonant frequency, we find that the MNM shows self-rectification, which is key for precisely timed neural stimulation (Fig. 2B). The red line shows a 200 μs window sliding average of voltage waveform, which reveals a 1.7 V bias voltage develops across the MNM but not across the ME film (Fig 2A) when a magnetic field is applied (see methods). These data also show that by pulsing the AC magnetic field we can generate a DC electrical stimulus. Figure 2A and 2B show the example of an arbitrary low-frequency pulse train using a 335 kHz carrier. Thus, we can generate a low-frequency electrical stimulus using the high-frequency magnetic carrier signal. This ability to demodulate the high-frequency carrier to a low-frequency electrical impulse is what enables us to generate a waveform that can directly activate voltage gated ion channels that would otherwise not respond to a high-frequency alternating electric field.

As a proof of principle, we demonstrate the capability of the MNM to restore a fast sensory motor reflex in an anesthetized rat. For this experiment we use the MNM to stimulate the sciatic nerve when we measure mechanical stimulation of the rat’s foot using a force sensor (Fig. 3A). Under awake or lightly sedated conditions, rats display a sensory reflex where the leg muscle contracts producing a leg kick when the foot is pinched. (Fig. S4) When rats are fully anesthetized this sensory response is inhibited and the lack of this sensory motor reflex is often used to check whether the animal is fully anesthetized. To test if the MNM could function as part of a neuroprosthetic capable of restoring sensory motor reflexes we programmed the magnetic field driver to stimulate the MNM at a 1 Hz frequency when a force sensor on the rat foot measures a mechanical stimulation. For these experiments, we placed a rat under anesthesia and surgically exposed the sciatic nerve (see methods). We then placed a MNM on the exposed nerve and applied a magnetic field with an external field generator at the skin’s surface, which is approximately 1 cm away from the nerve and the force sensor on the foot of the rat (see methods) (Fig. 3A). With the rat fully anesthetized we observed no EMG or leg kicks when we pinched the rat’s foot. However, when we activated the MNM with a remote magnetic field pulse (1.5 mT, 1 ms) in response to the force sensor, we observed restoration leg kicks EMG signals when we pinched the toe demonstrating that MNMs can act as part of a neuroprosthetic system (Fig. 3B). We found that the MNM was capable of reliable stimulation at frequencies of 1, 2, and 10 Hz when driven at the resonant frequency suggesting that these MNM could support a variety of neuroprosthetic applications that require millisecond timing. (Fig. 3C, Fig. S5). To confirm that the neural stimulation was the result of the self-rectifying MNMs and not the magnetic field stimulus, we detuned the magnetic field driver away from the resonance frequency by less than 5% and found that both the leg kicks and EMS responses vanished. (Fig. 3D) We also observed no leg kicks or EMG response when we stimulated an ME laminate at its resonance frequency (Fig 3E), confirming that the self-rectifying properties of the MNM is key to enabling rapid stimulation of neural activity.

As another example of how the precisely timed MNM stimulation can support neuroprosthetic applications we used the MNM to restore nerve conduction across a severed rat sciatic nerve (Fig. 4). As shown in Fig. 4A, we used a pulse generator to stimulate upstream of the severed nerve with a 1 mA, 1 ms pulse and used a nerve cuff to record the nerve potential at this proximal end of the severed nerve. We then used this voltage measured on the nerve cuff as a trigger for the magnetic field driver to activate an electrical pulse in the MNM on the distal end of the severed nerve. In this way we use the MNM to bridge the nerve gap allowing the electrical signal to continue across the severed nerve relying on the short latency of the MNM to avoid significant signal propagation delay. (Fig. 4A). We plot the applied and measured voltages in Fig. 4. We found that the MNMs successfully bridge the sciatic nerve gap and enable nerve signals to activate the distal muscle groups with latencies of less than 5 ms (Fig 4C). This response time is approximately 120-fold faster than any previously reported remote neural stimulation using magnetic materials (*6*). Unlike existing magnetic materials used for neural stimulation, we show high temporal control with our MNMs, demonstrated by both the short latency between pulse generator and the EMG response (4.06 +/− 0.209 ms) as well as the ability to drive neural responses at frequencies up to 10 Hz (Fig 4C, Fig S5). Furthermore, the latency between the pulse generator and the MNM across the nerve gap is only ~175 us. These fast latencies and associated higher stimulation bandwidths are important for many neurotherapeutic applications such as neuroprosthetics where sensory information needs to be quickly transmitted from a prosthetic (as demonstrated with the force sensor) or for bridging a signal propagation across a severed nerve.

**Fig. 4.**
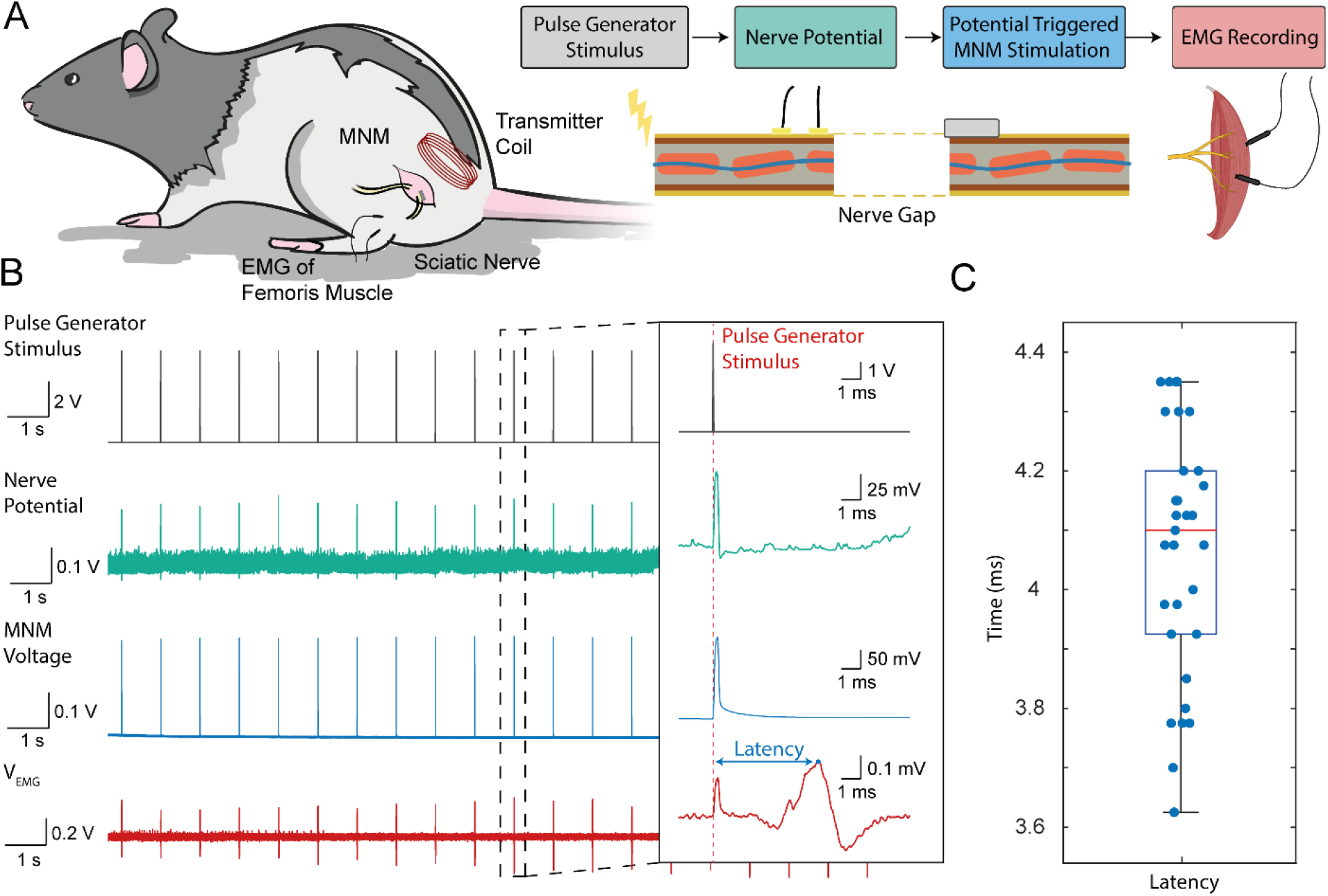
(A) Schematic showing how the MNM can support the restoration of nerve conduction across a severed sciatic nerve in vivo. The flow chart and corresponding nerve schematic shows the experimental setup to bridge the nerve gap. A compound nerve action potential is initiated by a pulse generator (PG) and the downstream nerve action potential (NAP) is detected at the proximal nerve end with cuff electrodes. The measurement of the NAP with the cuff electrodes then triggers the application of the magnetic field that drives the MNM in contact with the distal end of the nerve. This in turn stimulates a NAP on the distal side of the nerve thus bridging the nerve gap and activating a muscle response. (B) Voltage recordings on four channels corresponding to the PG stimulus, nerve potential, MNM voltage, and EMG voltage. The inset shows an expanded view of a single bridged nerve signal. (C) Boxplot of the latency of the EMG resulting from the initial PG stimulus across the nerve gap.

The ability to engineer nonlinear magnetoelectric effects not previously observed in ME materials is the key element that enables these magnetic metamaterials to achieve precisely timed stimulation of neural activity. An additional advantage is that we achieve this precise remote stimulation of neural activity without any genetic manipulation of the target tissue, which eases the path for potential clinical applications. Compared to other magnetic materials for neural interfaces that can be as small as 10 nm in diameter, the MNMs demonstrated here range in size from millimeters to centimeters in length. It remains an open question if MNM materials can be fabricated at a similar nanometer length scale and if they will work as efficiently for remote neural stimulation. Fortunately, the concept of MNMs presented relies on a nonlinear charge transport layer that is only 250 nm thick, and thus it may be possible to create nanoscale MNMs given recent advances in thin film ME fabrication technologies (*32*). Given this new metamaterial concept, more work is needed to develop theoretical treatments to describe the metamaterial properties. The MNM we demonstrate here relies on a nonlinear charge transport layer that affects the relationship between the electric displacement and electric field. This nonlinear effect is likely to be best represented as an effective nonlinear permittivity, but a theoretical description of MNMs is not fully developed here, and alternative electronic elements could be used to create MNMs with a variety of linear and non-linear effective dielectric properties. Additionally, other types of nonlinearities could be introduced by engineering different aspects of the magnetic-to-electric transduction pathway. For example, for strain-mediated magnetoelectric materials like those described here, it may be possible to engineer the mechanical properties of the laminate and introduce nonlinearities in the magnetoelectric coupling by drawing from work in acoustic metamaterials (*33*). A potential concern for chronic biotechnology applications is encapsulation of the MNM to avoid dampening the ME coefficient, and some materials like PZT layer which contains lead, may need to be substituted for more inert piezoelectric materials like AlN or PVDF. In summary we show that it is possible to rationally design metamaterials that convert magnetic fields to electric fields to overcome challenges in neurotechnology, but this framework for metamaterial design can likely be applied to diverse applications in sensing, electronics, and memory that rely on manipulating magnetic and electric fields in miniature materials.

## Methods

### Fabrication of ME

The ME material was fabricated by sandwiching a piezoelectric material between two sheets of a magnetostrictive material. A sheet of Metglas was first cleaned using Isopropyl Alcohol (IPA). The Hardman double bubble 2-part epoxy was then applied to the Metglas sheet and a sheet of lead zirconate titanate (PZT – 5A) was placed on top to form the composite material. This was left to dry for 5 minutes at room temperature. Another sheet of Metglas cleaned with IPA was bonded to this laminate, using the same epoxy to form a sandwich structure. The films were then cut to a desired size using a femtosecond laser cutter. Conductive Ag epoxy is used to connect 30 AWG wire to make electrical contact with the ME film so that the voltage could be tested. The same method was used to fabricate 2-layer ME materials where a sheet on Metglas was attached to a PZT sheet using the method described above (*16*). We characterized the two- and three-layer ME composites and found that they have qualitatively equivalent characteristics (Fig. S6A and Fig 1D-green dots).

### Fabrication of MNM

For fabrication of the MNM, a Metglas sheet was first plasma cleaned for 2 minutes. One side of the Metglas was then masked and on the other side 50 nm of Pt was deposited by sputtering using (AJA ATC Orion sputter, MA, USA) at 15 W in the presence of 30 sccm Argon for 20 minutes. Following this, 40nm of HfO_2_ was deposited as the passivating layer. Atomic Layer Deposition (Cambridge Ultratech Savannah 200) was used to get a conformal layer. Deposition was done at 150C using Tetrakisdimethylamido (TDMA) and H_2_O. On top of this, ALD was used to deposit 130nm of ZnO with the diethyl zinc (DEZ) precursor and H_2_O. This deposition was done at 100C, with precursor pulses of 0.015 seconds for DEZ and 0.015 seconds for H_2_O and a 45s reaction time between pulses to control the charge carrier concentration in this layer to achieve the self-rectification properties. The masking on the Metglas was then removed and the pristine side of this Metglas was attached using the non-conductive epoxy, to a laminate of Metglas and PZT fabricated using the same methods as those used to make the 2-layered ME material and was cut to an appropriate size. Silver epoxy was used as the metal contact to connect the ZnO to the bottom of the ME film using polyamide tape to insulate the remaining stack of materials. Figure 3 (left) shows the cross section of the entire material. Figure 3B (right) shows the false-colored SEM of the cross section of the deposited RET layers.

#### MNM Characterization

The fabricated RET stack follows the typical curves of a Schottky diode (Fig. S2A) with a turn-on voltage of 0.3V. We measured the I-V curves using a source meter (Keithley 2400). From the I-V curves, we calculated the rectification ratio. We obtained a rectification ratio of 95.38 at 10V. This was calculated by using the equation RR(V)=I(V)/I(-V). This is comparable with the rectification ratios of commercial silicon diodes which are between 105 and 108. We then fit the Shockley equation to our RET I-V curves as follows:

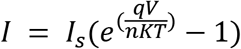

Where, V is the applied voltage, T=298K is the absolute temperature (room temperature), K is the Boltzmann constant and q is the charge of an electron.

By fitting the Shockley diode equation to the I-V curves we found that the RET had an ideality factor of 6.8, and a Saturation current Is=1.69 x10^-8^ which is comparable to other thin film diodes (*34*). These values clearly indicate that the RET layer forms a Schottky diode that modifies the effective permittivity within the MNM to provide the self-rectifying characteristics. We plotted the log I-log V curves (Fig S2B). The linear curve in the positive region and the non-linearity in the negative region shows that the electron transport in this material occurs through the SCLC mechanism and tunneling currents are restricted in the negative bias mode (*26*). To additionally verify the rectifying characteristics, we connected the MNM material in series with a signal generator (BK Precision 4052) and measured the output of the test circuit on an oscilloscope (Tektronix TBS1052B). We see a rectified output voltage for an input sine wave (Fig. S6B). To further verify that the ZnO/HfO_2_ combination and specific fabrication methodology results in the obtained rectification, we fabricated a structure with Al2O3 as the charge transport layer. We found no rectification as seen from the I-V curves (Fig. S6D).

### ME and MNM coupling coefficient

To calculate the ME and MNM coupling coefficients, we used a 3 mm × 2 mm ME and MNM films with a mechanical resonance frequency of 345KHz and 335KHz respectively. The films were wirelessly driven with a magnetic field at their respective resonant frequencies. We varied the amplitude of the AC magnetic field between 0 to 0.5mT and used an oscilloscope (Tektronix TBS1052B) to measure the output maximum and minimum voltage values, which correspond to the maximum and minimum applied magnetic field. The graph of measured output voltage and applied magnetic field is plotted in Fig 1D (bottom) for both ME and MNM. We then used a polynomial function to fit the measured curves to obtain the coupling coefficients for ME and MNM. We also did the same measurement for MNM between −1.5mT to 1.5mT, since we used a 1.5mT field for the in vivo experiments. We see that the MNM material saturates beyond a certain magnetic field strength, however the trend of the output voltage follows the nonlinearity introduced by the diode layer. The results are shown in supplementary Fig. S7C.

### ME and MNMH-I curves

To compare the characteristics of ME and MNM, and to verify that the nonlinear characteristics of MNM were a direct result of the addition of the RET, we first measured the I-V curves of the RET layer at the MNM operational frequency of 335 kHz. To do this, we used a signal generator (BK Precision 4052), connected to a separately fabricated RET layer with an external resistance of 470k ohm in series as shown in Fig S3A(inset). We varied the input voltage and measured the current across the load resistance to plot the RET I-V curves in Fig S3A. Then we fit the I-V curve to a 5th order polynomial that accurately captures the measured data. For the ME film, we mapped the applied magnetic field values to the film output voltage values using H-V curve (fig 1D) then we mapped these values to the output current of the ME film + RET using the 5th order polynomial of the RET IV curve. The ME H-V curve from Fig 1D was then plotted as a function of this fit. The curve obtained through this is plotted in Fig S3B. This curve represents the output current measured when the output voltage obtained from the ME material is applied as an input to the RET. We then measured the H-I curve of the MNM by measuring the current across a 470 kΩ resistance while applying a resonant magnetic field to the MNM. We compare the curves in Fig S3B and S3C in Fig 2C.

### Magnetic Field Transmitter

A custom magnetic field driver was built to drive our coils. The entire transmitter system included the use of an Arduino that was programmed through USB from a laptop. The microcontroller sent two signals to the driver to control the H-bridge in order to drive biphasic square pulses through the magnetic coil. The system included a 30-turn coil with an inner diameter of 17 mm, wrapped with 30 AWG wire. The driver is capable of supporting up to 1.25 MHz and 30 Amps of continuous current.

### In Vivo Rat Sciatic Nerve

All procedures carried out in this research complied with the National Institutes of Health standards and were approved by the Animal Care and Use Committee of Rice University (Protocol #IACUC-20-181). The MNMs were used to stimulate the sciatic peripheral nerve of Long Evans rat models. Successful stimulation of the rat hind legs by the MNMs were confirmed with recorded EMG and observable leg kicks. For the surgical procedure, the animals were placed under anesthesia with an induction chamber with 5% isoflurane in oxygen at a flow rate of 1-2 liters per minute. When the rat was unconscious and areflexic, confirmed with toe pinches, the animal was transferred to a 40°C heated pad with a nose cone to provide ~2% isoflurane. Meloxcam 2 mg/kg SC and Ethiqa XR SC 0.65 mg/kg were administered to the rat prior to shaving around the surgical site. The site was then sterilized using iodine swabs before a semi-circular incision was cut across the hip of the rat. The fascial plane between the gluteus maximus and anterior head of the bicep femoris was dissected to expose the peripheral nerve. Underlying connective tissue was carefully cut to isolate the sciatic nerve. The MNM is then placed directly onto the sciatic nerve and the magnetic coil is brought to ~1 cm of the material. EMG is recorded through two needle electrodes on the subplantar regions with a reference electrode connected to the main body of the rat. The recording system included a DAQ and dual bioamplifier (ADsystems) sampling at 1 kHz. The data was preliminarily observed through LabChart before exporting to Matlab 2017 and 2020 for processing. Upon completion of the experiments, the animals were immediately euthanized under proper guidelines.

For the experiments, the MNMs were made with 3 different dimensions-- sample 1: 10 × 5mm, sample 2: 5 × 3mm and sample 3: 3 × 2mm. The bottom Metglas layer was extended for the extra surface area to deposit electrodes on the same plane of the material. The three samples were used wireless stimulation of the rat sciatic nerve where the silver epoxy pads fabricated on the MNM surface were brought in contact with the sciatic nerve (Fig 3A inset) and the MNMs were wirelessly driven by magnetic fields at 1.5mT and a DC bias magnet to obtain maximum voltages (Fig S5C). For all three samples (Sample1: 100KHz, Sample 2: 242KHz, Sample 3:335Khz), we observed leg kicks and recorded the EMG response. Additional control data was obtained to verify that the MNMs only work at the mechanical resonant frequency of the material. For this, MNMs were driven off-resonance (Sample 1: 90KHz, Sample 2: 230KHz, Sample 3: 330 kHz). We observed no leg kicks and no EMG signal was obtained with off resonance stimulation (Fig 3B, middle). To calculate the response latency, we used MATLAB 2017 to quantify the time difference between the initial PG pulse and the first peak of the EMG response. These experiments were performed with MNMs of various sizes ranging between 3 mm and 10 mm in length with resonance frequencies varying between 100 to 365kHz. The latencies and EMG responses were similar to those evoked by direct electrical stimulation with wired electrodes (Fig. S6C)

For the severed nerve experiment, the sciatic was cut using surgical shears. Platinum stimulating electrodes were hooked up to the ADsystems pulse generator. A stimulating pulse of 1 mA with 1 ms pulse width was applied upstream of the proximal nerve ending. Platinum recording electrodes were then used to measure resulting potential at the severed proximal end of the nerve. This recorded voltage was then amplified and used as a trigger to initiate a single millisecond long magnetic field pulse that would power the MNM on the distal severed nerve ending. EMG electrodes were placed in the bicep femoris of the rat to record resulting muscle activity.

To test the on and off resonant controls using the ME laminates, polyamide was used to insulate one edge of the film before adding the conductive epoxy. One conductive epoxy electrode was placed in the center of the ME film while the other electrode on the reverse side was deposited in the center and wrapped around the edge of the insulated film so that both electrodes would reside on the same side and plane. The ME laminates were driven at 335 kHz on resonance and at 325 kHz for the off resonant control by bringing the resonant coil to the surface of the rat. This resulted in a recorded artifact pulse but no observable leg kick or recorded EMG. We further verified that the physiological response was indeed a function of the rectification and modulation of the high carrier frequency to the lower pulsed frequency. In this case, we powered Pt electrodes that were brought into contact with the sciatic nerve using a wired ME film (5 mm × 3 mm) driven with a 245 kHz magnetic field. When we pulsed our magnetic field at 1 Hz with a 1 ms pulse width, we observed no response. Upon adding a commercial diode in parallel with the ME material, we observed leg kicks as well as recorded EMG signals that corresponded with the applied stimulation pulses. (Fig. S7)

#### *In Vivo* experiment for sensory reflex restoration

The rats were induced under isoflurane with the same protocol mentioned above in the general rat procedure. A control experiment was run where the isoflurane concentration is reduced from the normal 2% to 0.5% to restore the toe pinch reflex response. A force sensor (Tekscan Flexiforce 101) was placed on the foot of the rat and used to visualize the toe pinch force and resulting EMG response was recorded from the femoris muscle of the rat (Fig. S4). The force sensor voltage read out saturates at 5V which is the voltage rail on the amplifier circuit. The isoflurane was then increased back to the usual 2% where the toe pinch test was executed and no EMG and muscle activation was observed, thus indicating a fully anesthetized animal. The same surgical procedure was then carried out to expose the sciatic nerve. A noninverting amplifier (TL061CP) was used to amplify the sensed voltage. The arduino teensy was then configured to initiate 1 Hz stimulation when the sensor voltage reached a threshold. Occasionally, there would not be a resulting MNM voltage upon toe pinch as can be seen in Fig. 3B, this is due to slight misalignment of the transmitter to the metamaterial and thus not inducing a high enough voltage.

## Acknowledgments

We would like to acknowledge Amanda Singer, Anne Tuppen, Matthew Parker, and Vishnu Nair for their useful discussions on magnetoelectrics. We would also like to acknowledge Edwin C. Lai for his assistance with rat nerve surgeries. We also thank the staff at the Shared Equipment Authority (SEA) at Rice, Tim Gilheart, Jing Guo, James Kerwin, Hua Guo for their assistance and excellent discussions.

## Funding

National Science Foundation ECCS-2023849 (GB, JCC, FA, JTR)

National Institutes of Health U18EB029353 (GB, JCC, FA, JTR)

## Author contributions

Conceptualization: GB, JCC, JTR
Methodology: GB, JCC
Investigation: GB, JCC, FA, AD
Data Curation: GB, JCC
Funding acquisition: JTR
Supervision: JTR
Writing – original draft: GBB, JCC, JTR
Writing – review & editing: GBB, JCC, FA, AD, JTR

## Competing interests

Authors declare that they have no competing interests.

## Data and materials availability

All data are available in the main text or the supplementary materials. Any unprocessed data and code used are available upon reasonable request.

## Supplemental

**Figure S1.**
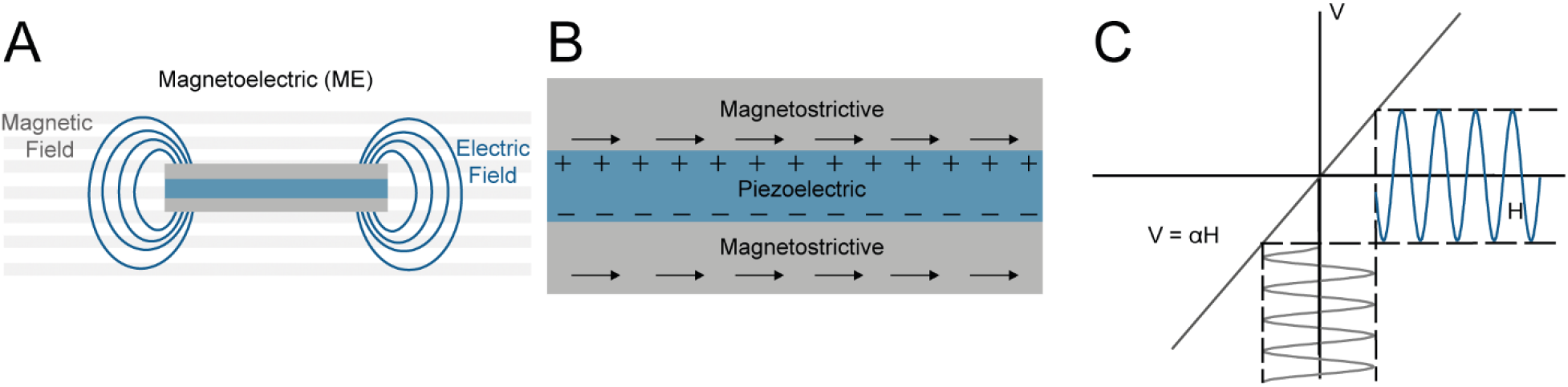
(A) Schematic of a magnetoelectric laminate. Upon application of a magnetic field, the magnetostrictive layer is mechanically deformed. The strain is then transferred to the piezoelectric layer generating an electric field. (B) Schematic of the ME layers which consists of a piezoelectric layer between two magnetostrictive layers. (C) Conceptual graph of H vs V curve which demonstrates a linear relationship. The inset shows the equivalent circuit model for the ME laminate.

**Figure S2.**
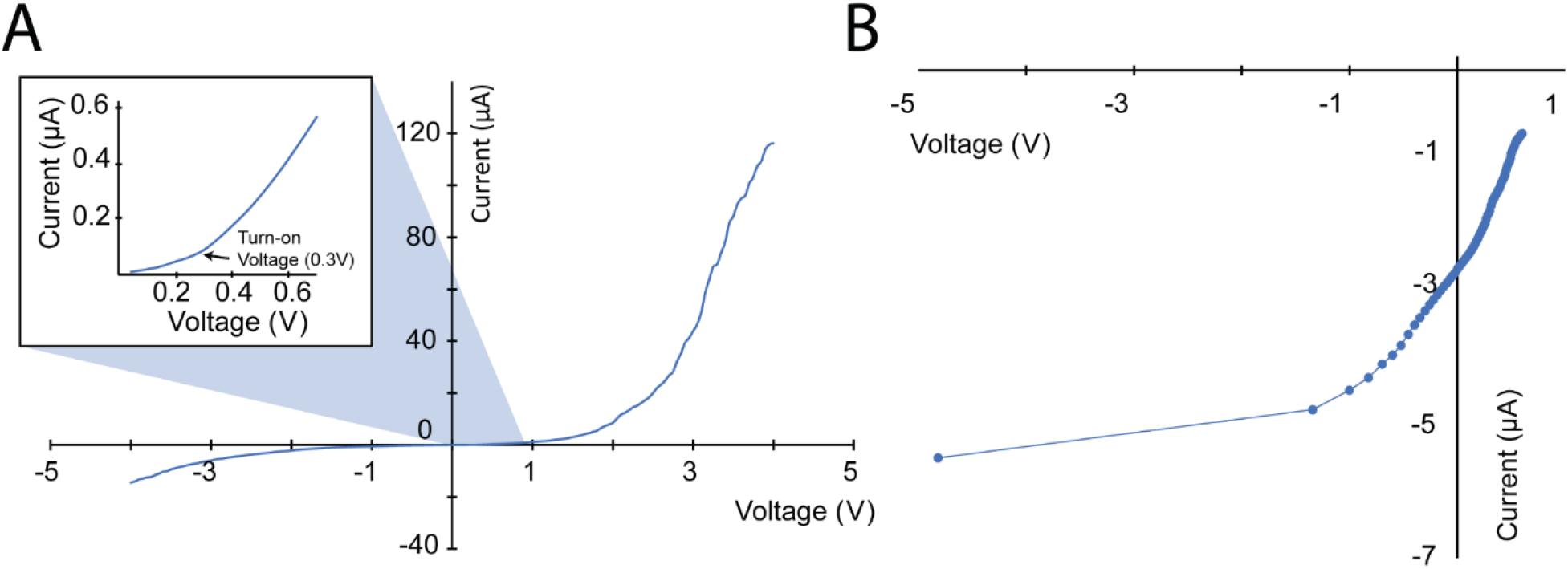
(A) I -V curve of MNM material. Inset: Zoomed in view, shows turn-on voltage ~0.3V. (B) Log I-log V curves for MNM material. The nonlinear characteristics in the negative region and linear characteristics in the positive region show charge transport through the SCLC mechanism. (*35*)

**Figure S3.**
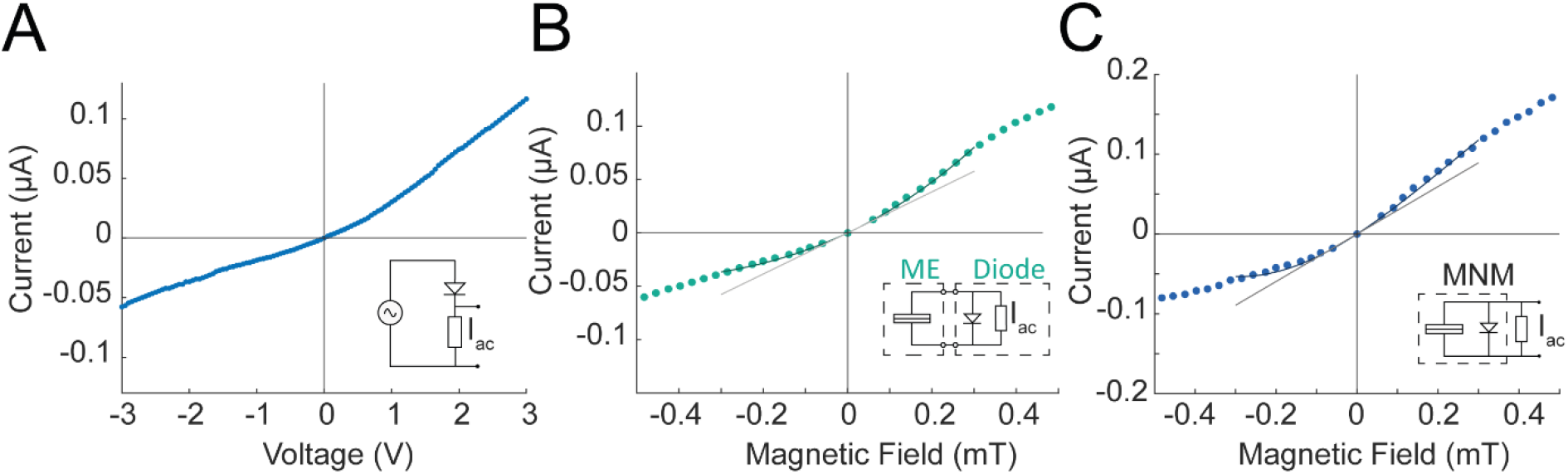
(A) The I-V curve of only the transport layers taken at 335 kHz with the data taken using the equivalent circuit model shown. (B) Mapping the linear ME voltage to the I-V curve of the charge transport layers yields the current vs applied magnetic field strength. The dark green 3rd order polynomial fit is plotted in contrast to a grey linear fit. (C) The MNM I-V characteristics measured with a load resistor also demonstrates a nonlinear function as demonstrated with the equivalent circuit model. The dark blue polynomial fit is plotted along with a grey trend line.

**Fig. S4.**
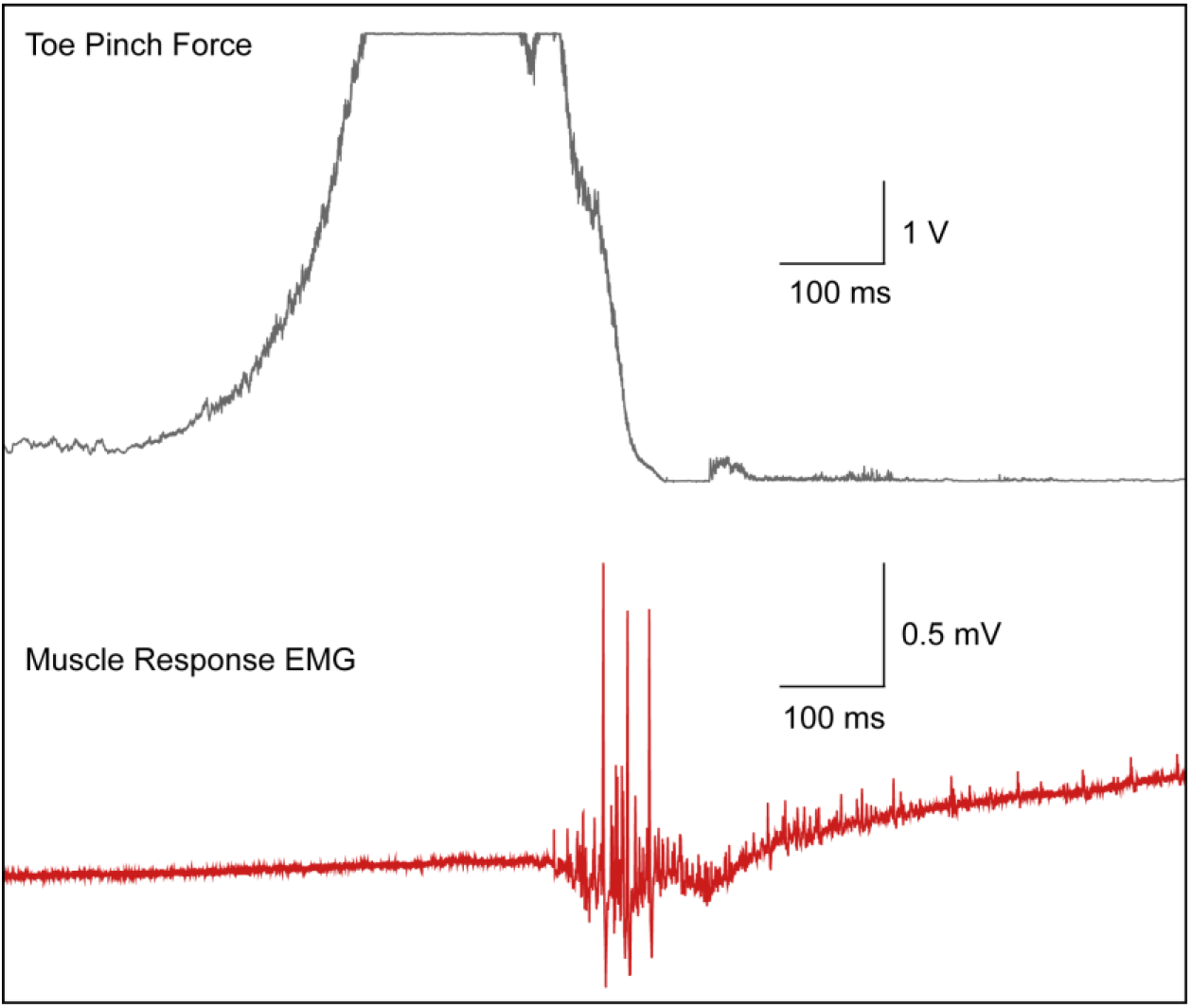
EMG response of a naive rat lightly sedated with 0.5% isoflurane to a toe pinch. (red) Force of toe pinch is also measured and shown with a sensor placed on the foot. (grey)

**Fig. S5:**
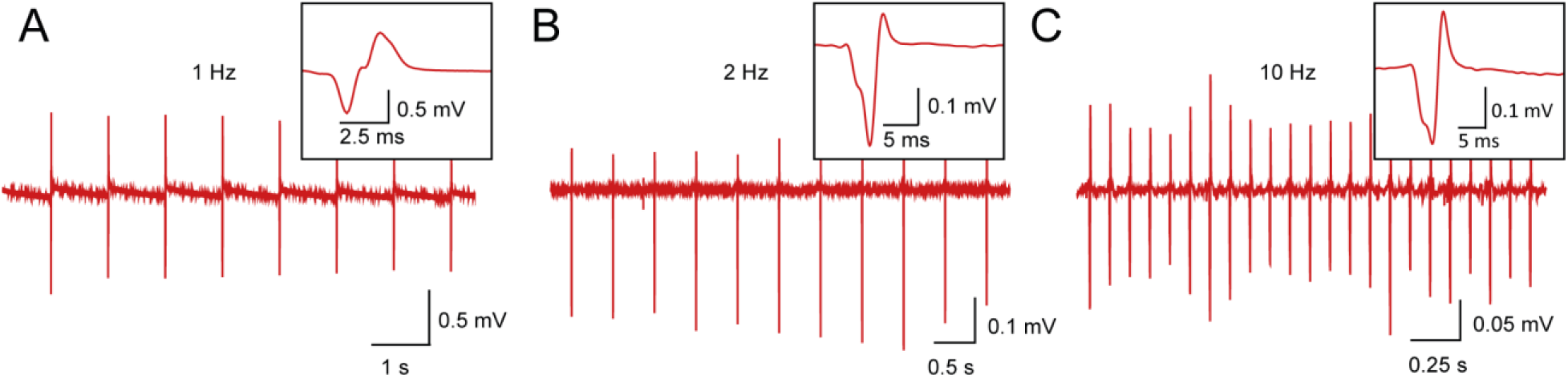
(A) EMG response to MNM stimulus at 1 Hz (B) EMG response to MNM stimulus at 2 Hz (C) EMG response to MNM stimulus at 10 Hz. EMG data presented in this figure are from 3 different rats.

**Figure S6.**
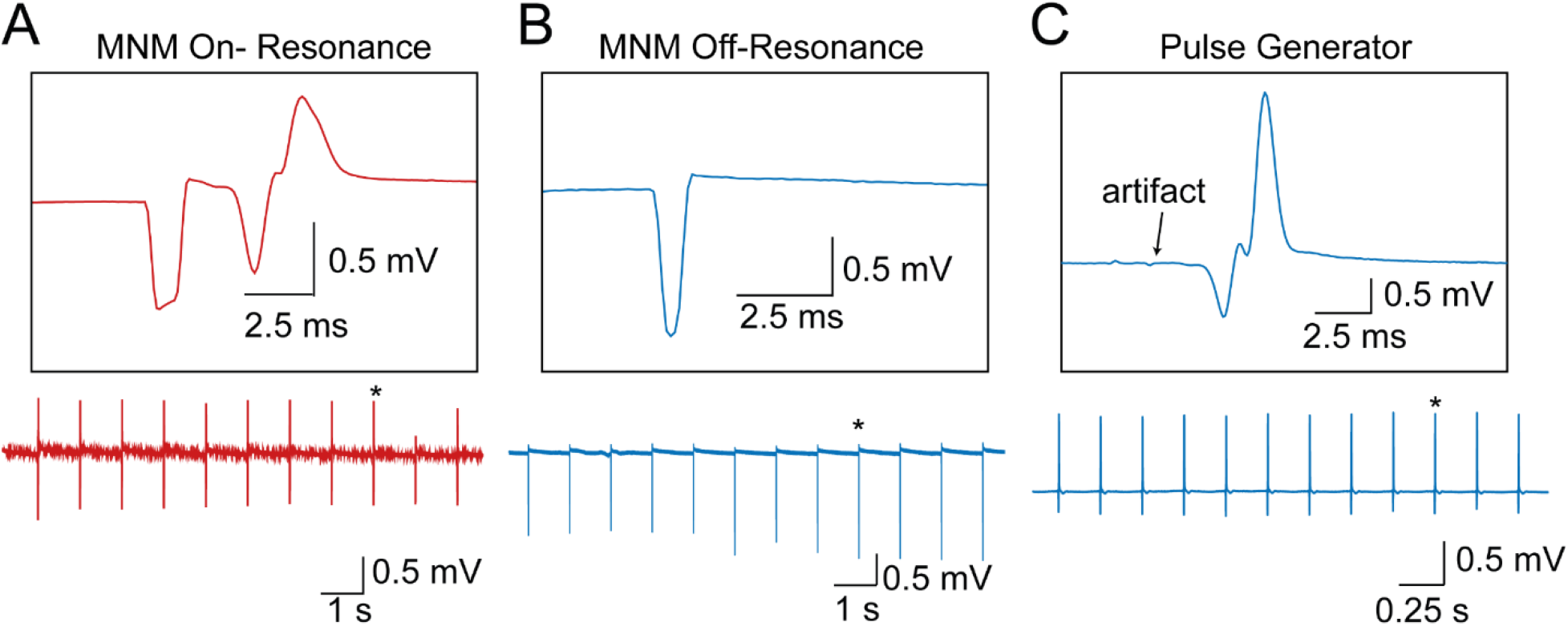
(A) EMG response including stimulus artifact with MNM stimulating with resonant frequency of 365 kHz at 1 Hz. Zoom in is of the waveform indicated by the asterisk in the lower plot. (B) Recorded electrical activity on the same EMG channel with an off-resonant magnetic field of 350 kHz. Waveform shows the stimulus artifact but no following EMG response. Zoom in is of the waveform indicated by the asterisk in the lower plot. (C) EMG response to a biphasic pulse with a 2 V amplitude, 1.5 ms pulse width from an external pulse generator.

**Figure S7.**
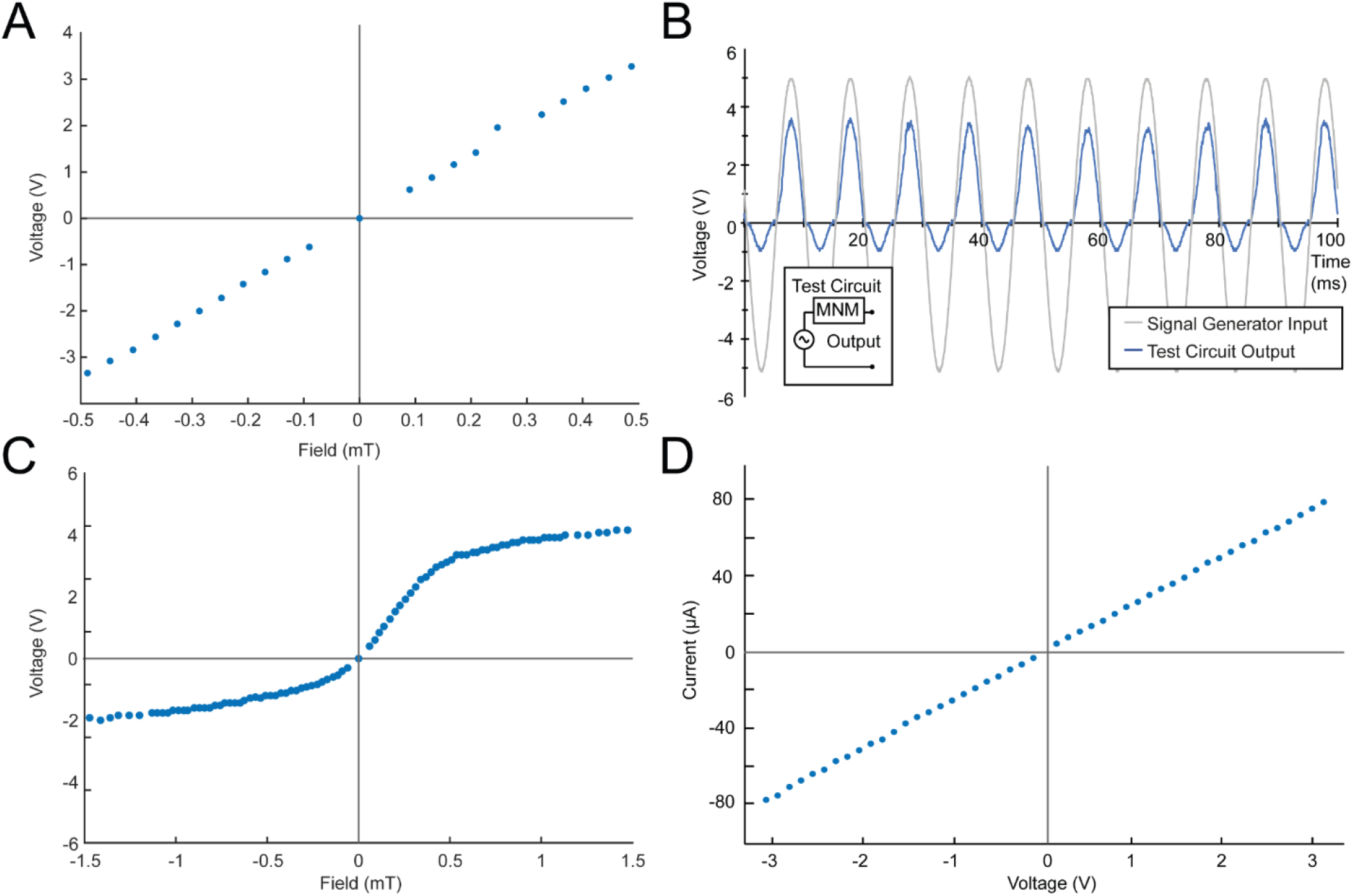
(A) Linear H-V curves from −0.5 to 0.5 mT for a three-layered laminate consisting of Metglas-PZT-Metglas. (B) Rectified output voltage obtained from an MNM connected in series with a signal generator, when a sine wave is applied as an input. Inset shows the test circuit. (C) H-V curve for the MNM from −1.5 to 1.5 mT field strengths. (D) I-V curve of a sham MNM laminate. 100 nm of Al2O3 is used to replace the 100 nm of ZnO in the charge transport layers, which results in a linear response.

**Figure S8.**
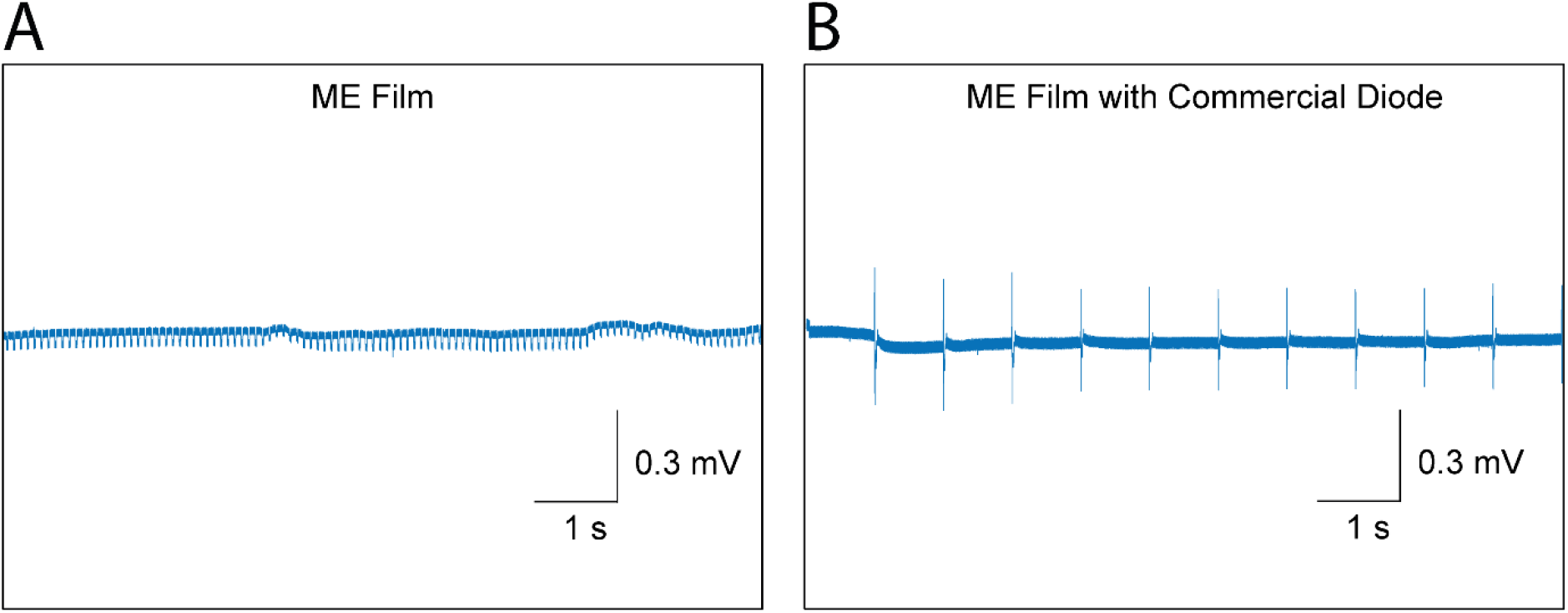
(A) Stimulation of the sciatic nerve with just an ME film at 345 kHz with no EMG response or leg kicks. (B) Stimulation of the sciatic nerve with the same ME film in parallel with a commercial diode resulting in leg kicks and an EMG response.

## Notes

### Competing Interest Statement

The authors have declared no competing interest.

### Summary of Updates

Fixed referencing errors, supplemental figure numbering errors, and updated author list.

## References

1. M. Hallett et al. Transcranial magnetic stimulation and the human brain. Nature, 406(6792), 147–150 (2000)

2. R. Chen et al., Wireless magnetothermal deep brain stimulation. Science. (2015)

3. H. Huang et al., Remote control of ion channels and neurons through magnetic-field heating of nanoparticles. Nature Nanotechnology. 5, 602–6 (2010).

4. R. Munshi, A. Pralle, Remote modulation of neuronal cells in the brain, Nature Materials vol. 20, p. 912–913 (2021)

5. J. Moon et al., Magnetothermal multiplexing for selective remote control of cell signaling. Adv. Funct. Mater. 30, 2000577.(2020)

6. C. Sebesta, et. al., “Sub-second multi-channel magnetic control of select neural circuits in behaving flies”, bioRxiv (2021)

7. S. Rao et al., Remotely controlled chemomagnetic modulation of targeted neural circuits. Nature Nanotechnology, 14(10), 967–973. (2019)

8. E.S. Boyden et al., Millisecond-timescale, genetically targeted optical control of neural activity. Nature Neuroscience, 8(9), 1263–1268. (2005)

9. D. Gregurec et al., Magnetic Vortex Nanodiscs enable remote magnetomechanical neural stimulation. ACS Nano, 14, 8036–8045, (2020)

10. J. Dobson et al., Remote control of cellular behaviour with magnetic nanoparticles. Nature Nanotechnology, 3(3), 139–143. (2008)

11. J. Lee et al., Non-contact long-range magnetic stimulation of mechanosensitive ion channels in freely moving animals. Nature Materials, 20(7), 1029–1036. (2021)

12. C. Tu et al., Mechanical-Resonance-Enhanced Thin-Film Magnetoelectric Heterostructures for Magnetometers, Mechanical Antennas, Tunable RF Inductors, and Filters, Materials (Basel). 12(14):2259 (2019)

13. B. Hutcheona, Y. Yaromb, “Resonance, oscillation and the intrinsic frequency preferences of neurons”, Trends in Neurosciences, 23,5,1, pp. 216–222 (2000)

14. T. Nguyen et al., In Vivo Wireless Brain Stimulation via Non-invasive and Targeted Delivery of Magnetoelectric Nanoparticles. Neurotherapeutics (2021).

15. K. Kozielski et al., Nonresonant powering of injectable nanoelectrodes enables wireless deep brain stimulation in freely moving mice. Science Advances, 7 (2021)

16. A. Singer et al., Magnetoelectric materials for miniature, wireless neural stimulation at therapeutic frequencies, Neuron 107, 631–643 (2020)

17. Z. Yu, et. al., “MagNI: a magnetoelectrically powered and controlled wireless neurostimulating implant, IEEE Transactions on Biomedical Circuits and Systems (TBioCAS), 14,6, pp. 1241–1252 (2020)

18. F. Alrashdan et al. Wearable wireless power systems for ‘ME-BIT’ magnetoelectric-powered bio implants, Journal of Neural Engineering, 18 (2021)

19. J.C. Chen et al. Wireless endovascular nerve stimulation with a millimeter-sized magnetoelectric implant, bioRxiv (2021)

20. L.Y. Fetisov et al. Nonlinear magnetoelectric effects at high magnetic field amplitudes in composite multiferroics. Journal of Physics D: Applied Physics, 51(15). (2018)

21. Y.K. Fetisov et al. Frequency dependence of magnetoelectric voltage for a multilayer ferrite-piezoelectric structure with finite conductivity, Integrated Ferroelectrics, 106:1, 23–28 (2009)

22. E. Almeida et al. Nonlinear metamaterials for holography. Nature Communications, 7, 1–7 (2016)

23. N.I. Zheludev, Y.S. Kivshar, From metamaterials to metadevices. Nature Materials, 11(11), 917–924 (2012)

24. L.W. Martin et al. Multiferroics and magnetoelectrics: Thin films and nanostructures. Journal of Physics Condensed Matter, 20(43). (2008)

25. T. D. Cuong, et al. Giant magnetoelectric effects in serial-parallel connected Metglas/PZT arrays with magnetostrictively homogeneous laminates, Journal of Science: Advanced Materials and Devices, 5,3, pp. 354–360 (2020)

26. Y. Park, et. al., Unidirectional oxide hetero-interface thin-film diode, Appl. Phys. Lett. 107, 143506 (2015)

27. T. A. Krajewski, et. al. Hafnium dioxide as a passivating layer and diffusive barrier in ZnO/Ag Schottky junctions obtained by atomic layer deposition, Appl. Phys. Lett. 98, 263502 (2011)

28. L. J. Brillson and Y. Lu, ZnO Schottky barriers and Ohmic contacts, Journal of Applied Physics 109, 121301 (2011)

29. S. Mondal, et. al., Preparation of ZnO Film on p-Si and I-V Characteristics of p-Si/n-ZnO, Materials Research. 16(1): 94–99 (2013)

30. E. Guziewicz, et. al., ALD grown zinc oxide with controllable electrical properties, Semicond. Sci. Technol. 27, 074011, (2012)

31. S. Jeon, et. al., Structural and electrical properties of ZnO thin films deposited by atomic layer deposition at low temperatures, Journal of The Electrochemical Society, 155,10, H738–H743, (2008)

32. M. Zaeimbashi et al. Ultra-compact dual-band smart NEMS magnetoelectric antennas for simultaneous wireless energy harvesting and magnetic field sensing. Nature Communications, 12(1) (2021)

33. S.A. Cummer et al. Controlling sound with acoustic metamaterials. Nature Reviews Materials, 1(16001) (2016)

34. N. Ahmed, Numerical analysis of transport properties of ZnO based Schottky diode, Phys. Scr. 96 (2021)

35. F.C. Chiu, A Review on Conduction Mechanisms in Dielectric Films, Adv. Mat. Sci & Eng., 578168 (2014)

